# Indolergic receptors of the elephant mosquito *Toxorhynchites amboinensis*

**DOI:** 10.1101/513044

**Authors:** Dekel Amir, Esther Yakir, Jonathan D. Bohbot

**Affiliations:** Department of Entomology, The Hebrew University of Jerusalem, Rehovot 76100, Israel

**Keywords:** Odorant receptor, indole, skatole, mosquito, *Toxorhynchites amboinensis*, indolergic receptor

## Abstract

The conservation of the mosquito indolergic receptors across the Culicinae and Anophelinae mosquito lineages, which spans 200 million years of evolution, is a testament to the central role of indolic compounds in the biology of these insects. Indole and skatole have been associated with the detection of oviposition sites and animal hosts. To evaluate the potential ecological role of these two compounds, we have used a pharmacological approach to characterize homologs of the indolergic receptors *Or2* and O*r10* in the non-hematophagous elephant mosquito *Toxorhynchites amboinensis*. We provide evidence that both receptors are narrowly tuned to indole and skatole like their counterparts from hematophagous mosquitoes. These findings indicate that indole and skatole are operating in a non-animal-host seeking context in *Toxorhynchites* and underscore the importance of understanding their roles in hematophagous mosquitoes.

## 1. Introduction

It is well-established that resource-locating mosquito behaviors are mainly mediated by olfactory signals (Takken and Knols, 1999; Zwiebel and Takken, 2004). However, only a few relevant animal/plant hosts and oviposition odorants have been identified (Davis and Bowen, 1994). In this regard, the ecological roles of indole and its close analog skatole (3-methylindole), two nitrogen-containing aromatic compounds, are complex (Figure 1A). Indole and skatole alone or as a mixture, have been proposed to act as oviposition attractants in *Aedes aegypti* (Baak-Baak et al., 2013), *Culex* spp. (Beehler et al., 1994; Blackwell et al., 1993; Du and Millar, 1999; Mboera and Takken, 1999; Millar et al., 1994; 1992; Mordue et al., 1992) and *Anopheles gambiae* (Lindh et al., 2008). These two compounds, products of the metabolic activity of the microflora, are present in significant amount in mammalian waste products (Garner et al., 2007; Yokoyama and Carlson, 1979), human skin (Bernier et al., 2000; 2002) and human sweat (Meijerink et al., 2000), which indicate they may also act as animal-host attractants (Cork, 1996). Indole is a ubiquitous component of flower scents of many plant families (Knudsen et al., 2006) and it has been identified from host plants of *Ae. aegypti* and *An. gambiae* (Nyasembe et al., 2018).

**Figure 1.**
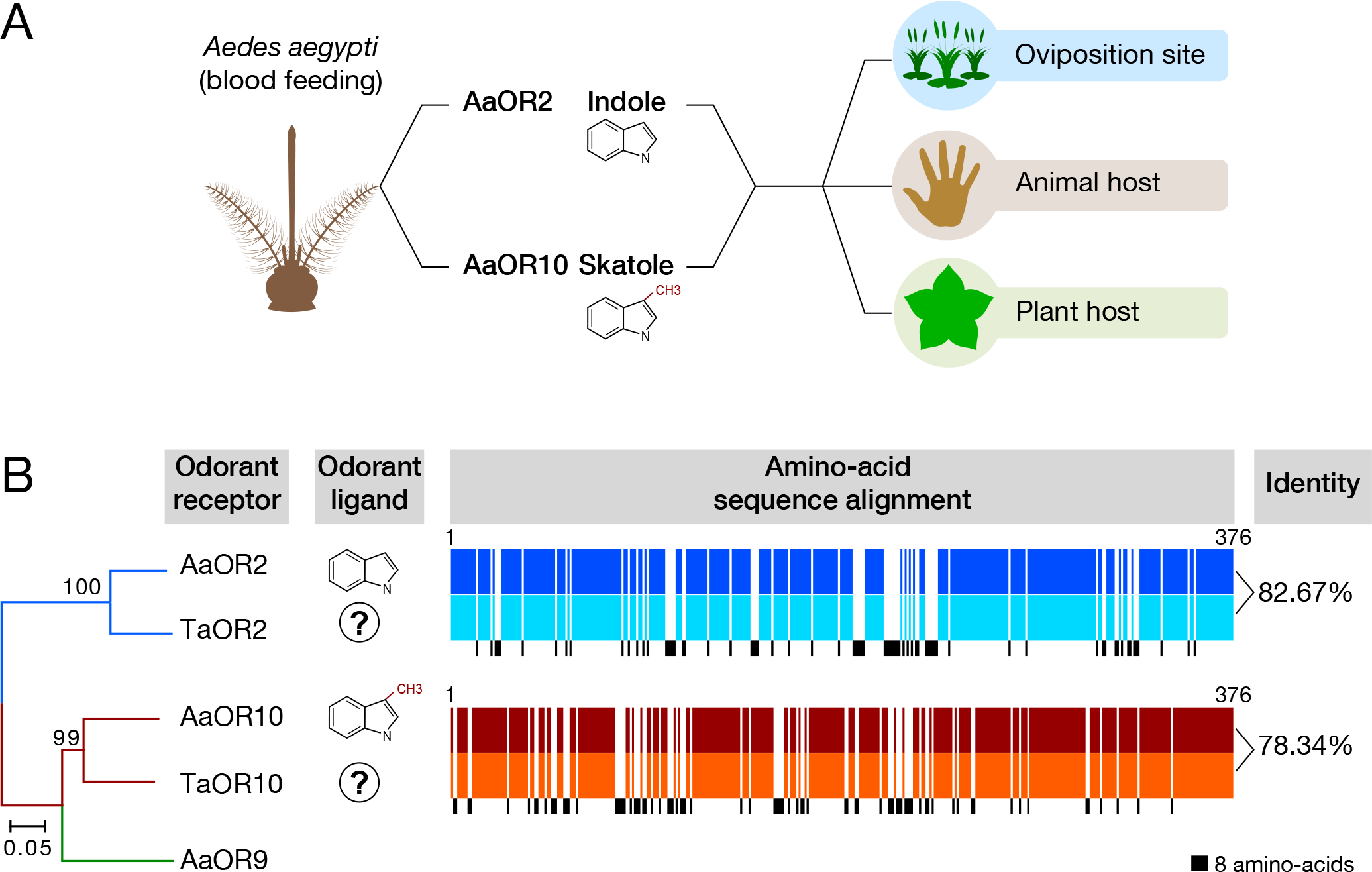
Indolergic receptors may operate in multiple ecological contexts. (A) Genes encoding the *Aedes aegypti* odorant receptors 2 (AaOR2) and 10 (AaOR10) proteins are expressed in the adult antennae and are respectively activated by indole and skatole. These two compounds have been linked to oviposition site selection and animal-host seeking in *Aedes aegypti*. Indole is also a major component of flower scents and may play a role in plant-host attraction. (B) Phylogenetic analysis of the candidate indolergic receptor from *Toxorhynchites amboinensis*. Amino-acid identity between OR2 and OR10 are colored in blue and red, respectively. Amino-acid differences are shown in black. AaOR9 was used as an outgroup.

Both indole and skatole are electrophysiologically active compounds detected by the antennae of *Ae. aegypti* (Siju et al., 2010), *C. quinquefasciatus* (Du and Millar, 1999) and *An. gambiae* (Blackwell and Johnson, 2000; Meijerink et al., 2000; 2001; Qiu et al., 2006). Two highly conserved odorant receptors (ORs), OR2 and OR10, are highly sensitive and selective towards indole and skatole, respectively in *Ae. aegypti* (Bohbot et al., 2011), *C. quinquefasciatus* (Hughes et al., 2010; Pelletier et al., 2010) and *An. gambiae* (Carey et al., 2010; Wang et al., 2010). The *Or2* and *Or10* genes are expressed in the antennae of adult male and female mosquitoes, while *Or2* is only expressed in the antennae of larvae along with a third paralog named *Or9*. The functional conservation of these receptors across the two mosquito subfamilies in both sexes and in larvae suggest that indole and skatole play important and multiple roles in the biology of these insects (Figure 1A) (Cork, 1996; Nyasembe et al., 2018).

Our objective was to explore the role of these two compounds in the context of animal-host selection by functionally characterizing candidate homologs of the indole and skatole receptor genes in the non-hematophagous elephant mosquito *Toxorhynchites amboinensis*. Lack of functional conservation would argue the case for a role of the OR2-indole and OR10-skatole pairs in animal-host seeking in hematophagous insects. Our study shows that *T. amboinensis* OR2 (TaOR2) and OR10 (TaOR10) share high sequence identity with their *Aedes* counterparts in support of a highly conserved role in mosquitoes outside animal-host seeking. We provide pharmacological evidence that the elephant mosquito TaOR2 and TaOR10 are indole and skatole receptors operating in a non-animal host context, including oviposition site selection and/or plant-host-seeking.

## 2. Materials and Methods

### 2.1. Cloning *TaOr* genes and sequence analyses

Cloning and sequencing of *TaORco* was described elsewhere (Dekel et al., 2016a). *TaOr2* and *TaOr10* were custom-synthesized (Bio Basic Inc., Markham Ontario, Canada), subcloned into into the pENTR^TM^ vector using the Gateway^R^ directional cloning system (Invitrogen Corp., Carlsbad, CA, USA) and subcloned into the *Xenopus laevis* expression destination vector pSP64t RFA. Plasmid purification was carried out using the The ZR Plasmid Miniprep^TM^-Classic (Zymo Research, Irvine, CA, USA) and sequenced by Macrogen Europe (Amsterdam, the Netherland). DNA and amino-acid sequences for *TaOr2*, *TaOr10* and *TaORco* have previously been published (Zhou et al., 2014) and can be accessed at Figshare (https://figshare.com/articles/Transcriptome_assembly_of_T_ambionsis/2182684/2, last accessed on Dec 26, 2018).

Amino-acid sequence alignments (Supplementary Figure 1) were executed using MAFFT version 7 (Nakamura et al., n.d.). Phylogenetic analysis was performed using the neighbor-joining statistical function and 10,000 Bootstrap replications of the MEGA 7 software (Kumar et al., 2016).

### 2.2. Chemical reagents

For establishing the tuning curve, we used 30 odorants, including 19 compounds from Sigma-Aldrich (Milwaukee, WI, USA), including 1-hepten-3-ol (CAS 4938-52-7), 3-methylbutanol (CAS 123-51-3), *E*-2-hexen-1-al (CAS 6728-26-3), heptaldehyde (CAS 111-71-7), octanal (CAS 124-13-0)c,etpatreop(CylA-aS 109-60-4), 3-octanone (CAS 106-68-3), 6-methyl-5-hepten-2-one (CAS 110-93-0), 2,4,5-trimethylthiazole (CAS 13623-11-5), diallyl-sulfide (CAS 2179-57-9), benzaldehyde (CAS 100-52-7), indole (CAS 83-34-1), histamine (CAS 51-45-6), (+)-limonene oxide (CAS 203719-54-4), geranyl-acetate (CAS 105-87-3), (+)- fenchone (CAS 4695-62-9), 2-oxopentanoic acid (CAS 1821-02-9), (±)-1-octen-3-ol (CAS 3391-86-4) and 3-methylindole (CAS 83-34-1); 7 compounds from Merck (Darmstadt, Germany), including methyloctanoate (CAS 111-11-5), ethyl-hexanoate (CAS 123-66-0), 2-heptanone (CAS 110-43-0), dimethyl-sulfide (CAS 2179-57-9), tryptamine (CAS 61-54-1), octanoic-acid (CAS 124-07-2) and D-glucuronolactone (CAS 32449-92-6); 2 compounds from Acros Organics (Thermo Fisher Scientific, Waltham, MA, USA), including methyl salicylate (CAS 119-36-8) and octopamine (CAS 770-05-8); and 2 compounds from Alfa-Aesar (Ward Hill, MA, USA), including L-lactic acid (CAS 79-33-4) and δ-Decalactone (CAS 705-86-2).

### 2.3. Two-electrode voltage clamp electrophysiological recording of *Xenopus* oocytes expressing TaOR2, TaOR10 and TaORco

The methodologies and protocols used in this study have been described elsewhere (Bohbot and Dickens, 2009). Briefly, *TaOr2*, *TaOr10* and *TaORco* cRNA were synthesized using linearized pSP64tRFA expression vectors as template for in vitro transcription according to the instructions of the mMESSAGE mMACHINE® SP6 Transcription Kit (ThermoFisher Scientific). Stage V-VI oocytes were manually separated and enzymatically defolliculated using a 1 mg/mL collagenase (Sigma-Aldrich, Milwaukee, WI, USA) solution (calcium-free ND96 buffer, [pH 7.6]) for 40-50 min at 18 °C. Oocytes were then successively washed in calcium-free ND96 and gentamycin-supplemented (10 mg/mL, Sigma-Aldrich, Milwaukee, WI, USA) calcium-free ND96. Oocytes were then washed and incubated in ND96 buffer supplemented with cal- cium (0.1 M), 5% heat-inactivated horse serum (ThermoFisher Scientific), 50 mg/ml tetracycline (Carl Roth GmbH), 100 mg/ml streptomycin (Sigma-Aldrich, Milwaukee, WI, USA) and 550 mg/ml sodium pyruvate (Sigma-Aldrich, Milwaukee, WI, USA) for four to five days. Oocytes were injected with 27.6 nL (27.6 ng of each cRNA) of RNA using the Nanoliter 2010 injector (World Precision Instruments, Inc., Sarasota, FL, USA). Odorant-induced currents of oocytes expressing *TaOr2/10* and *TaORco* were recorded using the two microelectrode voltage-clamp technique (TEVC). The OC-725C oocyte clamp (Warner Instruments, LLC, Hamden, CT, USA) maintained a −80 mV holding potential.

For the establishment of concentration-response curves, oocytes were exposed to to indole or skatole alone 10^−10^ M to 10^−4^ M). Data acquisition and analysis were carried out with the Digidata 1550 A digitizer and pCLAMP10 software (Molecular Devices, Sunnyvale, CA, USA).

The tuning curve was generated using a panel of 30 odorants including indole, skatole and other compounds known to elicit physiological or behavioral responses in mosquitoes. All chemicals used were administered at 90 nM, which approximates the EC_50_ of indole and skatole. All the data analyses were performed using GraphPad Prism 7 (GraphPad Software Inc., La Jolla, CA, USA).

## 3. Results

### 3.1. TaOR2 and TaOR10 are highly conserved

The antennae of *Toxorhynchites amboinensis* express two highly conserved indolergic receptor homologs (TaOR2 & TaOR10). Multiple sequence alignments and phylogenic analyses show that they share about 80% amino-acid identity with their *Aedes* counterparts (Figure 1b). TaOR2 and AaOR2 encode 376 amino-acid proteins sharing 82.67% overall sequence identity. TaOR10 and AaOR10 encode 375 amino-acid proteins, which share 78.34% amino-acid identity. The OR10 alignment requires only one gap to be introduced (Supplementary Figure 1). Amino-acid divergence is evenly distributed throughout the peptide sequence. Both TaOR2 and TaOR10 grouped with their *Aedes* counterparts supported by bootstrap support with values above 95%.

### 3.2. TaOR2 and TaOR10 are highly sensitive to indolic compounds

Our next question was to investigate whether the observed sequence conservation determined functional orthology. To do so we expressed both receptors in the frog *Xenopus* oocyte system for pharmacological characterization.

Because *Aedes* and *Anopheles* OR2 are highly sensitive to indole (Bohbot et al., 2011; Wang et al., 2010), we challenged TaOR2 with ten-fold increasing concentrations of this odorant and skatole, a methylated analog of the former. Like its *Aedes* counterparts, TaOR2 proved to be a challenging receptor to express in *Xenopus* oocytes as it consistently generates comparatively lower currents than other mosquito ORs such as OR10 or OR8 (Figure 2A). The resulting concentration-response relationships derived from the maximum amplitudes elicited by each tested concentrations provided EC_50_ values of 88 nM for indole and 1,380 nM for skatole (Figure 2B). Indole was about 15 times more potent than skatole and the indole dynamic concentration range occurred in the nanomolar range.

**Figure 2.**
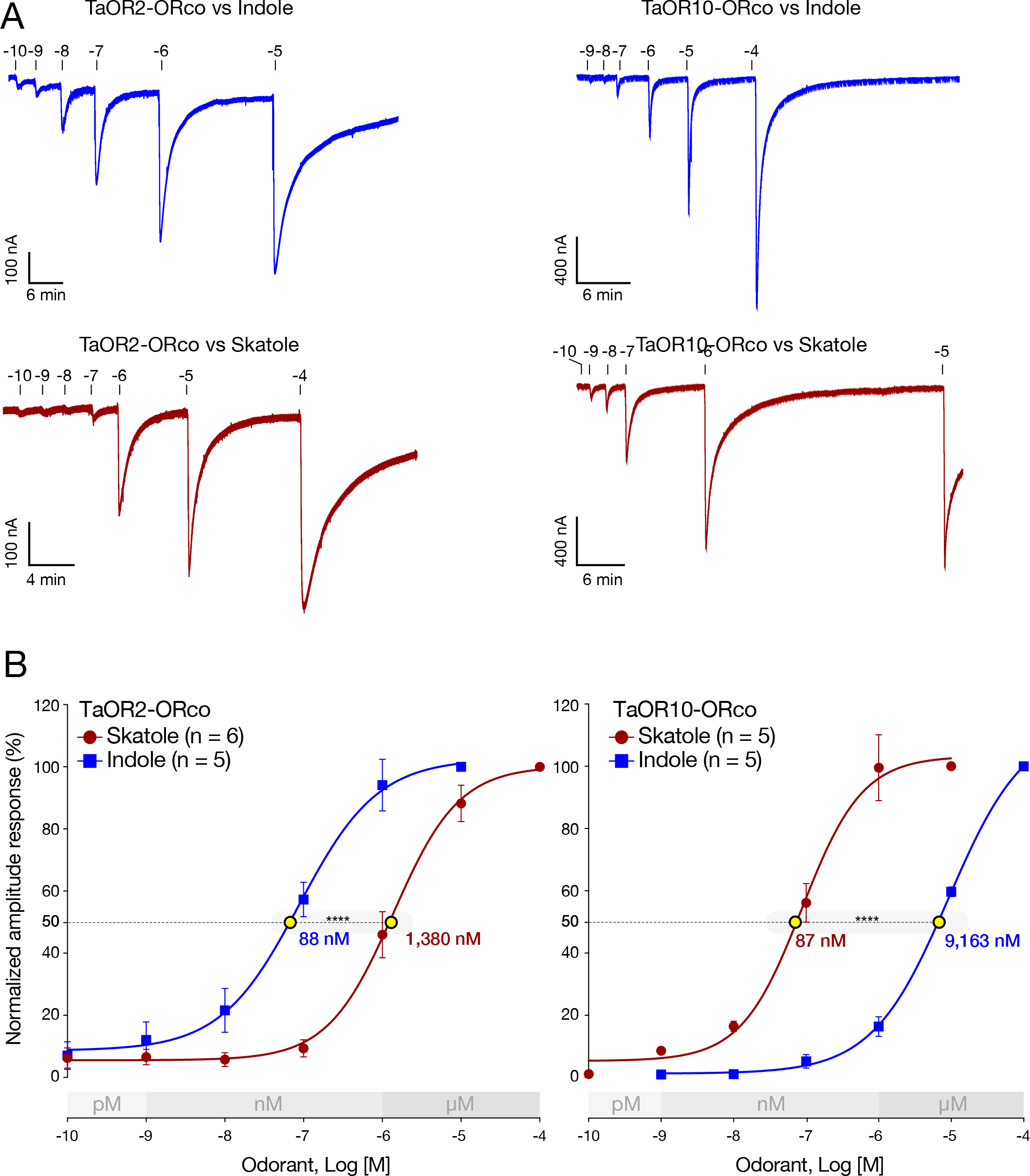
TaOR2 and TaOR10 are highly sensitive to indole and skatole, respectively. (A) Representative current traces elicited by increasing concentrations of indole and skatole recorded from *Xenopus* oocytes co-expressing the *TaOR2 or TaOr10* and *TaORco* receptor complexes. (B) Concentration-response relationships of TaOR2+TaORco and TaOR10+TaORco elicited by indole (blue curve) and skatole (red curve). Responses were normalized to the maximum amplitude response. Extrapolated EC_50_ values are shown with yellow circles. Lower and upper EC_50_ values (standard error) are in upper case. Asterisks represent statistically significant differences of the OR responses (one-way ANOVA followed by Tukey’s multiple comparisons post test; ****P < 0.0001). Odorant concentrations were plotted on a logarithmic scale. Each point represents the mean and error bars indicate s.e.m.

We applied the same approach to TaOR10 informed by the sensitivity of AaOR10 towards skatole (Bohbot and Dickens, 2012). Indeed, AaOR10 dynamically responds to skatole in the nM range while it is much less sensitive to indole (mM range). TaOR10 consistently generated robust and larger currents than the TaOR2 paralog (Figure 2A). The relative potency of indole and skatole was however reversed with TaOR10 being 105 times more sensitive to skatole (EC_50_ = 87 nM) than to indole (EC_50_ = 9,163 nM) (Figure 2B).

### 3.3. TaOR2 and TaOR10 are narrowly tuned to indole and skatole, respectively

We further tested the odorant selectivity of these two receptors using a panel of 30 compounds belonging to diverse chemical classes, including alcohols, aldehydes, esthers, ketones, sulfur compounds, aromatics, amines, terpenes, carboxylic acids and lactones (Dekel et al., 2016b) (Figure 3B). In order to avoid receptor adaptation, antagonist effects and technical artefacts such as broad molecular receptive ranges (Bohbot and Pitts, 2015) associated with high chemical concentrations (Bohbot and Pitts, 2015), the screens were carried out at low 90 nM odorant concentration, which nearly corresponds to the EC_50_ values of the TaOR2-indole and TaOR10-skatole pairs. We controlled for possible position effects by administering odorant sets in reverse order (Supplementary Figure 2). At this concentration, indole and skatole elicited the strongest responses for their respective cognate receptors (Figure 3A). We did not observe any modulation of receptor activity in response to the cognate ligands at the end of the recording sessions.

**Figure 3.**
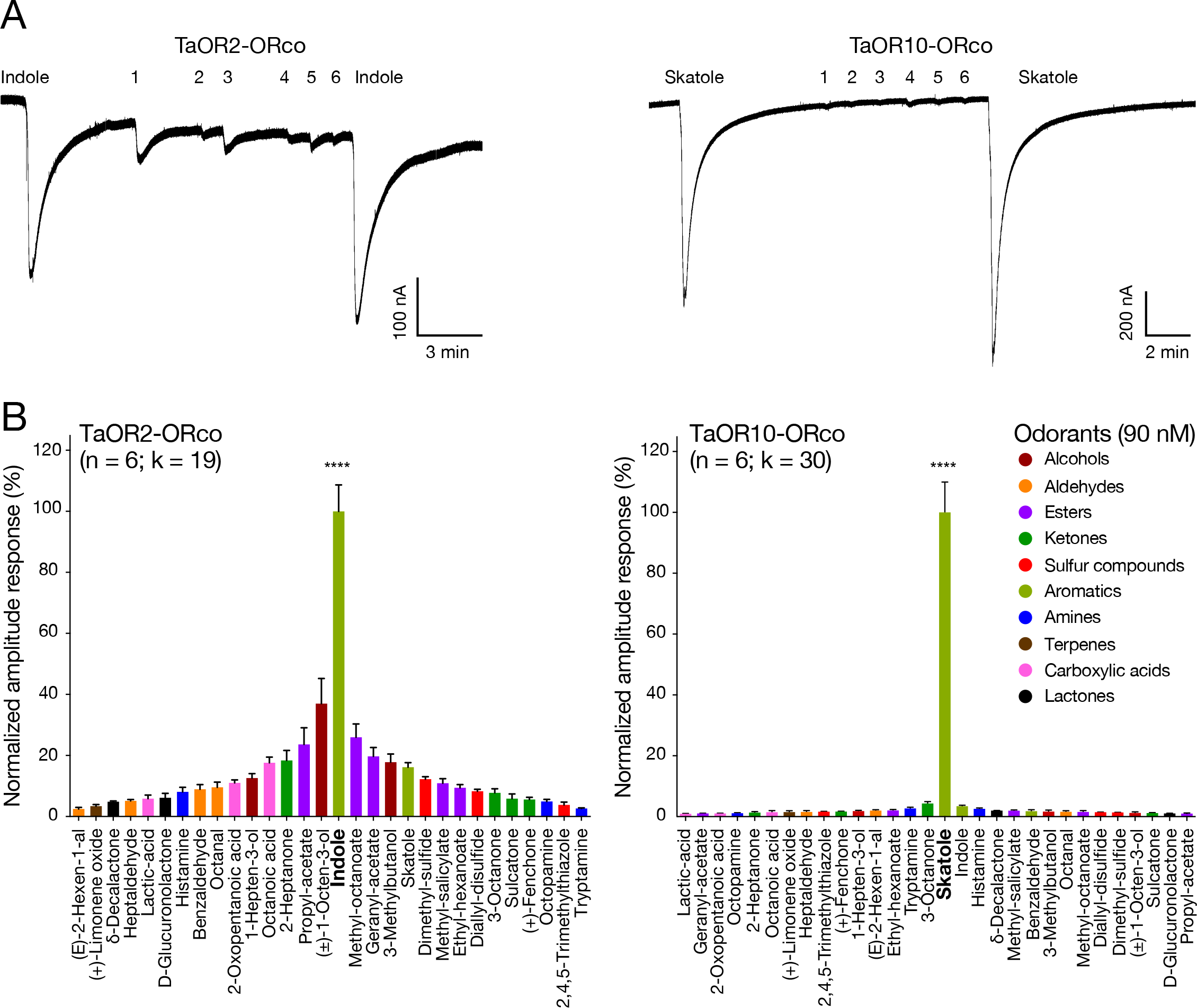
TaOR2 and TaOR10 are narrowly tuned. (A) Representative current trace elicited by indole, skatole, (±)-1-octen-3-ol (1), heptaldehyde (2), propyl-acetate (3), 3-octanone (4), benzaldehyde (5) and octopamine (6) recorded from *Xenopus* oocytes co-expressing the *TaOr2* or *TaOr10* and the *TaORco* receptor complexes. (B) TaOR2 and TaOR10 are narrowly tuned (k, kurtosis value) to indole and skatole, respectively (one-way ANOVA followed by Dunnet post-test; ****P < 0.0001). Mean responses (± s.e.m., n = 6) to 90 nM of 30 odorants were normalized to indole or skatole.

Overall, TaOR2 exhibited a medium response profile with a kurtosis value of 19 (Figure 3B). TaOR10 was narrowly tuned to skatole with a maximal kurtosis value of 30. These findings confirm the selectivity of these two receptors for skatole and indole, respectively. The activation of TaOR2 elicited by indole was 2.7 times greater than the next most potent odorant ((±)-1-octen-3-ol) and 3.8 times greater than the third most active odorant (methyl-octanoate). The activation of TaOR10 by skatole was 23.1 times higher than the next most activating odorant (3-octanone) and 29 times greater than indole.

## 4. Discussion

It is unclear whether indole and skatole act as oviposition or as animal-host (Cork, 1996; Millar et al., 1992). To complicate the matter, these benzenoid compounds participate to the flower scent of plants (Jürgens et al., 2010; Smith and Meeuse, 1966)1 and are preferred by Ae. aegypti in the context of plant-host attraction (Nyasembe et al., 2018). To explore the ecological role of indole and skatole in the biology of blood-feeding mosquitoes, i.e., to determine whether these receptors may operate in a non-animal host seeking context, we have pharmacologically tested the responses of the OR2 and OR10 homologs from the strict-nectar feeding mosquito *T. amboinensis*. Our criteria for determining functional orthology included both pharmacological sensitivity and selectivity properties. On both accounts, we provide evidence that TaOR2 and TaOR10 are functional orthologs of their counterparts from blood-feeding mosquito species, as they are highly sensitive (nanomolar range EC_50_ values) and narrowly tuned (kurtosis values) to indole and skatole, respectively.

While, the biological significance between these two receptors in terms of tuning breadth remains unresolved, the nanomolar level sensitivity of these receptors do suggest cognate relationships between these pairs. TaOR2 and TaOR10 exhibit high-level of sensitivity (nM range) among *T. amboinensis* ORs. For comparison, the EC_50_ value for (*R*)-1-octen-3-ol in relation to TaOR8 is 401 nM (Dekel et al., 2016a), about 4 times higher than TaOR2-indole and TaOR10-skatole interactions. The high sensitivity of TaOR2 and TaOR10 towards indole and skatole underscores their fundamental ecological significance.

TaOR10 exhibited outstanding specificity towards skatole, second to OR8 towards (*R*)-1-octen-3-ol (Bohbot and Dickens, 2009; Dekel et al., 2016a). Using a panel of 29 compounds identical to the ones tested here but excluding skatole, we find that the kurtosis value (k = 29) for TaOR8 was maximal akin to TaOR10. (*R*)-1-octen-3-ol was 30.3 times more potent than the next most activating odorant. Such ligand specificity is suggestive of the ecological significance of this odorant. Additionally, such a high degree of specificity may reflect an adaptation for high signal to noise ratio at the peripheral level (Lu et al., 2007).

We have provided evidence that TaOR2 and TaOR10 share the same function as their counterparts from the blood-feeding mosquitoes *Ae. aegypti*, *An. gambiae* (Bohbot et al., 2011; Bohbot and Dickens, 2012; Wang et al., 2010) and *Culex quinquefasciatus* (Hughes et al., 2010; Pelletier et al., 2010). In *Toxorhynchites* sp., these receptors may mediate oviposition site selection (Collins and Blackwell, 2002) and exhibit sensitive physiological responses (Collins and Blackwell, 1998). Our findings exclude the role of indole and skatole in animal-host seeking as far as *T. amboinensis* is concerned and underscore the need to decipher the role(s) of these compounds in blood-feeding mosquitoes using detailed behavioral studies. The unusual sequence and functional conservations of OR2 and OR10 during mosquito evolution reflect the importance of indole and skatole to mosquito ecology and behavior. Indeed, not only do adult detect indole but the larva antennae of *Ae. aegypti* also express *Or2*, suggesting a separate ecological role of this compound in aquatic environments. The prevalence and abundance of indole and skatole present us with a challenge that is to understand their potential role in foraging, mate searching, habitat finding and oviposition site seeking. In addition, odorants can elicit different activities from different mosquito species (Xu et al., 2015), which means that indole and skatole may operate in different contexts within and between species.

The conservation and central role of the mosquito-specific *Or2* and *Or10* genes may be leveraged for the development of future mosquito control agents, including receptor (agonists and antagonists) and behavioral modulators (repellents and attractants). However, comprehensive behavioral studies are wanting to develop such tools for vector population control and personal protection.

## Supplemental information

**Supplementary Figure 1.** (A) Amino-acid sequence alignment of OR2 and OR10 between *Toxorhynchites amboinensis* and *Aedes aegypti*. (B) Percentage of amino-acid sequence identity.

**Supplemental Figure 2.** The order of odorant administration does not affect the relative receptor activity. Representative current traces elicited by 90 nM of indole, skatole, (±)-1-octen-3-ol (1), heptaldehyde (2), propyl-acetate (3), 3-octanone (4), benzaldehyde (5) and octopamine (6) recorded from *Xenopus* oocytes co-expressing the *TaOr2* and the *TaORco* receptor complex.

## Acknowledgements

The authors They are grateful to Prof. Eitan Reuveny and Dr. Izhar Karbat from the Weizmann Institue of Science for their help with the frog oocytes. This work was supported by the by the Israel Science Foundation [grant number 1990/16].

## References

Baak-Baak, C.M., Rodríguez-Ramírez, A.D., García-Rejón, J.E., Ríos-Delgado, S., Torres-Estrada, J.L., 2013. Development and laboratory evaluation of chemically-based baited ovitrap for the monitoring of Aedes aegypti. J Vector Ecol 38, 175–181. doi:10.1111/j.1948-7134.2013.12024.x

Beehler, J.W., Beehler, J., Millar, J., Millar, J.G., Mulla, M.S., Mulla, M., 1994. Field evaluation of synthetic compounds mediating oviposition in Culex mosquitoes (Diptera: Culicidae). J Chem Ecol 20, 281–291

Bernier, U., Kline, D., Barnard, D., Schreck, C., Yost, R., 2000. Analysis of human skin emanations by gas chromatography/mass spectrometry. 2. Identification of volatile compounds that are candidate attractants for yellow fever mosquito (Aedes aegypti). Analytical Chemistry A 72: 747–756.

Bernier, U.R., Kline, D.L., Schreck, C.E., Yost, R.A., Barnard, D.R., 2002. Chemical analysis of human skin emanations: comparison of volatiles from humans that differ in attraction of Aedes aegypti (Diptera: Culicidae). J Am Mosq Control Assoc 18, 186–195.

Blackwell, A., Johnson, S., 2000. Electrophysiological investigation of larval water and potential oviposition chemo-attractants for Anopheles gambiae s.s. Ann Trop Med Parasitol 94, 389–398.

Blackwell, A., Mordue, A., Hansson, B., 1993. A behavioural and electrophysiological study of oviposition cues for Culex quinquefasciatus. Physiological Entomology 18, 343–348.

Bohbot, J.D., Dickens, J.C., 2012. Odorant receptor modulation: Ternary paradigm for mode of action of insect repellents. Neuropharmacology 62, 2086–2095. doi:10.1016/j.neuropharm.2012.01.004

Bohbot, J.D., Dickens, J.C., 2009. Characterization of an enantioselective odorant receptor in the yellow fever mosquito Aedes aegypti. PLoS ONE 4, e7032. doi:10.1371/journal.pone.0007032

Bohbot, J.D., Jones, P.L., Wang, G., Pitts, R.J., Pask, G.M., Zwiebel, L.J., 2011. Conservation of indole responsive odorant receptors in mosquitoes reveals an ancient olfactory trait. Chem Senses 36, 149–160. doi:10.1093/chemse/bjq105

Bohbot, J.D., Pitts, R.J., 2015. The narrowing olfactory landscape of insect odorant receptors. Frontiers in Ecology and Evolution 3, 39.

Carey, A.F., Wang, G., Su, C.-Y., Zwiebel, L.J., Carlson, J.R., 2010. Odorant reception in the malaria mosquito Anopheles gambiae. Nature 464, 66–71. doi:10.1038/nature08834

Collins, L., Blackwell, A., 2002. Olfactory cues for oviposition behavior in Toxorhynchites moctezuma and Toxorhynchites amboinensis (Diptera: Culicidae). J Med Entomol 39, 121–126

Collins, L.E., Blackwell, A., 1998. Electroantennogram studies of potential oviposition attractants for Toxorhynchites moctezuma and T. amboinensis mosquitoes. Physiological Entomology 23, 214–219.

Cork, A., 1996. Olfactory basis of host location by mosquitoes and other haematophagous Diptera., in: Olfaction in Mosquito Host Interactions. Ciba Foundation Symposium 200, pp. 71–88.

Davis, E.E., Bowen, M.F., 1994. Sensory physiological basis for attraction in mosquitoes. J Am Mosq Control Assoc 10, 316–325.

Dekel, A., Pitts, R.J., Yakir, E., Bohbot, J.D., 2016a. Evolutionarily conserved odorant receptor function questions ecological context of octenol role in mosquitoes. Sci Rep 6, 37330. doi:10.1038/srep37330

Dekel, A., Pitts, R.J., Yakir, E., Bohbot, J.D., 2016b. Evolutionarily conserved odorant receptor function questions ecological context of octenol role in mosquitoes. Sci Rep 6, 86. doi:10.1038/srep37330

Du, Y., Millar, J., 1999. Electroantennogram and oviposition bioassay responses of Culex quinquefasciatus and Culex tarsalis (Diptera: Culicidae) to chemicals in odors from Bermuda grass infusions. J Med Entomol 36, 158–166.

Garner, C.E., Smith, S., de Lacy Costello, B., White, P., Spencer, R., Probert, C.S.J., Ratcliffe, N.M., 2007. Volatile organic compounds from feces and their potential for diagnosis of gastrointestinal disease. FASEB J 21, 1675–1688. doi:10.1096/fj.06-6927.com

Hughes, D.T., Pelletier, J., Luetje, C.W., Leal, W.S., 2010. Odorant receptor from the southern house mosquito narrowly tuned to the oviposition attractant skatole. J Chem Ecol 36, 797–800. doi:10.1007/s10886-010-9828-9

Jürgens, A., Dotterl, S., Liede-Schumann, S., Meve, U., 2010. Floral scent composition in early diverging taxa of Asclepiadoideae, and Secamonoideae (Apocynaceae). South African Journal of Botany 76, 749–761. doi:10.1016/j.sajb.2010.08.013

Knudsen, J.T., Eriksson, R., Gershenzon, J., Ståhl, B., 2006. Diversity and Distribution of Floral Scent. The Botanical Review 72, 1–120. doi:10.1663/0006-8101(2006)72[1:DADOFS]2.0.CO;2

Kumar, S., Stecher, G., Tamura, K., 2016. MEGA7: Molecular evolutionary genetics analysis version 7.0 for bigger datasets. Molecular biology and evolution 33, 1870–1874. doi:10.1093/molbev/msw054

Lindh, J.M., Kannaste, A., Knols, B.G.J., Faye, I., Borg-Karlson, A.-K., 2008. Oviposition responses of anopheles gambiae s.s. (diptera: culicidae) and identification of volatiles from bacteria-containing solutions. J Med Entomol 45, 1039–1049. doi:10.1093/jmedent/45.6.1039

Lu, T., Qiu, Y.T., Wang, G., Kwon, J.Y., Rutzler, M., Kwon, H.-W., Pitts, R.J., Van Loon, J.J.A., Takken, W., Carlson, J.R., Zwiebel, L.J., 2007. Odor coding in the maxillary palp of the malaria vector mosquito Anopheles gambiae. Curr Biol 17, 1533–1544. doi:10.1016/j.cub.2007.07.062

Mboera, L., Takken, W., 1999. Odour-mediated host preference of Culex quinquefasciatus in Tanzania. Entomologia Experimentalis et Applicata 96:167-175 92, 83–88.

Meijerink, J., Braks, M., Brack, A., Adam, W., Dekker, T., Posthumus, M., Beek, T., Loon, J., 2000. Identification of olfactory stimulants for Anopheles gambiae from human sweat samples. Journal of Chemical Ecology 26(6) 1367–1382.

Meijerink, J., Braks, M.A., Van Loon, J.J., 2001. Olfactory receptors on the antennae of the malaria mosquito Anopheles gambiae are sensitive to ammonia and other sweat-borne components. J Insect Physiol 47, 455–464.

Millar, J.G., Chaney, J.D., Beehler, J.W., Mulla, M.S., 1994. Interaction of the Culex quinquefasciatus egg raft pheromone with a natural chemical associated with oviposition sites. J Am Mosq Control Assoc 10, 374–379.

Millar, J.G., Chaney, J.D., Mulla, M.S., 1992. Identification of oviposition attractants for Culex quinquefasciatus from fermented Bermuda grass infusions. J Am Mosq Control Assoc 8, 11–17.

Mordue, A.J., Blackwell, A., Hansson, B.S., Wadhams, L.J., Pickett, J.A., 1992. Behavioural and electrophysiological evaluation of oviposition attractants for Culex quinquefasciatus say (Diptera: Culicidae). Cell Mol Life Sci 48, 1109–1111. doi:10.1007/BF01947999

Nakamura, T., Yamada, K.D., Tomii, K., Katoh, K., 2018, n.d. Parallelization of MAFFT for large-scale multiple sequence alignments. Bioinformatics 34, 2490–2492.

Nyasembe, V.O., Tchouassi, D.P., Pirk, C.W.W., Sole, C.L., Torto, B., 2018. Host plant forensics and olfactory-based detection in Afro-tropical mosquito disease vectors. PLoS Negl Trop Dis 12, e0006185. doi:10.1371/journal.pntd.0006185

Pelletier, J., Hughes, D.T., Luetje, C.W., Leal, W.S., 2010. An odorant receptor from the southern house mosquito Culex pipiens quinquefasciatus sensitive to oviposition attractants. PLoS ONE 5, e10090. doi:10.1371/journal.pone.0010090

Qiu, Y.T., Van Loon, J.J.A., Takken, W., Meijerink, J., Smid, H.M., 2006. Olfactory Coding in Antennal Neurons of the Malaria Mosquito, Anopheles gambiae. Chem Senses 31, 845–863. doi:10.1093/chemse/bjl027

Siju, K.P., Hill, S.R., Hansson, B.S., Ignell, R., 2010. Influence of blood meal on the responsiveness of olfactory receptor neurons in antennal sensilla trichodea of the yellow fever mosquito, Aedes aegypti. J Insect Physiol 56, 659–665. doi:10.1016/j.jinsphys.2010.02.002

Smith, B.N., Meeuse, B.J., 1966. Production of volatile amines and skatole at anthesis in some arum lily species. Plant Physiol 41, 343–347.

Takken, W., Knols, B.G., 1999. Odor-mediated behavior of Afrotropical malaria mosquitoes. Annu Rev Entomol 44, 131–157. doi:10.1146/annurev.ento.44.1.131

Wang, G., Carey, A.F., Carlson, J.R., Zwiebel, L.J., 2010. Molecular basis of odor coding in the malaria vector mosquito Anopheles gambiae. Proc Natl Acad Sci USA 107, 4418–4423. doi:10.1073/pnas.0913392107

Xu, P., Zhu, F., Buss, G.K., Leal, W.S., 2015. 1-Octen-3-ol the attractant that repels. F1000Res 4, 156. doi:10.12688/f1000research.6646.1

Yokoyama, M.T., Carlson, J.R., 1979. Microbial metabolites of tryptophan in the intestinal tract with special reference to skatole. Am J Clin Nutr 32, 173–178. doi:10.1093/ajcn/32.1.173

Zhou, X., Rinker, D.C., Pitts, R.J., Rokas, A., Zwiebel, L.J., 2014. Divergent and conserved elements comprise the chemoreceptive repertoire of the non-blood feeding mosquito Toxorhynchites amboinensis. Genome Biol Evol 6, 2883–2896. doi:10.1093/gbe/evu231

Zwiebel, L.J., Takken, W., 2004. Olfactory regulation of mosquito–host interactions. Insect Biochemistry and Molecular Biology 34, 645–652. doi:10.1016/j.ibmb.2004.03.017

