## Supplementary Figure 1 for "Indolergic receptors of the elephant mosquito *Toxorhynchites amboinensis*"

# A

**TaOR2 vs AaOR2**

```
TaOR2           MLIENCPIINVNAKVWLFWGYIRQPKWYSYLLGCIPVTVLNVFQFMNLFHIIASGTGNMN
AaOR2           MLIENCPIINVNVKVWLFWAYLRKPKWYSYLLGCVPVTVLNVFQFMNLFHVIASGSGDMN
                ************.******.*:*:**********:***************:****:*:**

TaOR2           KIIIDGYFTVLYFNLVLRTSFLMTNRRKFEDFLQGVADEYAVLETEGEMRPLMEQLTRRA
AaOR2           KIIIDGYFTVLYFNLVLRTSFLMGNRGKFETFLEGIADEYAVLEKQNDIRPLLDQLTRRA
                *********************** ** *** **:*:********.:.::***::******

TaOR2           RILSNSNLWLGAFISVCFVTYPLFSQENVLPYGVYAPGVDVHATPIYEIVFVLQVYLTFP
AaOR2           RILSKSNLWLGAFISACFVTYPLFSPDSGLPYGVYIPGVDVHASPIYEIVFVLQIYLTFP
                ****:**********.********* :. ****** *******:**********:*****

TaOR2           ACCMYIPFTSFYCTCTLFGLIRIAALKRSLKQLQKFSGSQRTLQAKMKECFEYHKGIIKY
AaOR2           ACCMYIPFSSFYCTCALFGLVRIAALKRSLEKIHEYNTSPRSLFARIKECLQYHKDIIKY
                ********:******:****:*********::::::. * *:* *::***::***.****

TaOR2           VSELNELVTYIFLLEFLSFGMMLCALLFLLSTSNQLAQMLMIGSYIFMILSQMYALYWHS
AaOR2           VSDLNELVTYIFLLELLSFGMMLCALLFLLSISNQLAQMVMIGSYIFMILSQMYALYWHS
                **:************:*************** *******:********************

TaOR2           NEVREQSLEIGDSL-YNAGWLDFNRSIKKEMIMLIARAQRPLAIKVGNVYPMTLEMFQTL
AaOR2           NEVREQSLEIGDSLYYNSAWLDFDNSVKKKIILMLARAQRPLAIKIGNVYPMTLEMFQSL
                ************** **:.****:.*:**::*:::**********:************:*

TaOR2           LNVSYSYFTLLRRVYN
AaOR2           LNASYSYFTLLRRVYN
                **.*************
```

**TaOR10 vs AaOR10**

```
TaOR10          M-DILDCPLISVNARVWRFWSFVLKHDYMRYISIIPVSVMNVFMFADLYQAWGNMDVLII
AaOR10          MASILDCPIVSVNARVWRFWSFVLKHDAMRYISIIPVTVMTFFMFTDLCRSWGNIQELII
                * .*****::***************** *********:**..***:** ::***:: ***

TaOR10          NAYFAVLYFNAVLRTLILVHNHETYEGFLGRIASVYLEISLIDDDVVQKLVRDYTKRARM
AaOR10          KAYFAVLYFNAVLRTLILVKDRKLYENFMQGISNVYFEISHIDDHKIQSLLKSYTVRARM
                :******************:::: **.*:  *:.**:*** ***. :*.*::.** ****

TaOR10          LSISNLALGAIISTCFVVYPLFTGQRGLPYGMFIPGINNYESPQYEIFYVTQAVLTFPGC
AaOR10          LSISNLALGAIISTCFVVYPIFTGERGLPYGMFIPGLDSFRSPHYEIIYIVQVVLTFPGC
                ********************:***:***********::.:.**:***:*:.*.*******

TaOR10          CMYIPFTSFFATSTLFAMVQIKTLQRQLRTFKEGSIKGGDIDELKAKIVKFVEHHKRIIT
AaOR10          CMYIPFTSFFASTTLFGLVQIKTLQRQLQTFKD-NINSQDKEKVKAKVVKLIEDHKRIIT
                ***********::***.:**********:***: .*:. * :::***:**::*.******

TaOR10          YVSEMNDLVTYICLLEFLSFGMMLCALLFLLNIIENQAQIIIVAAYIFMIISQIFAFYWH
AaOR10          YVSELNSLVTYICFVEFLSFGMMLCALLFLLNVIENHAQIVIVAAYIFMIISQIFAFYWH
                ****:*.******::*****************:***:***:*******************

TaOR10          ANEVREESMGIAEAAYCGPWVELDDSIKKMLLLITLRAQQPLEITVGNVYTMTLEMFQSL
AaOR10          ANEVREESMNLAEAAYSGPWVELDNSIKKKLLLIILRAQQPLEITVGNVYPMTLEMFQSL
                *********.:*****.*******:**** **** ***************.*********

TaOR10          LNASYSYFTLLRRVYN
AaOR10          LNASYSYFTLLRRVYN
                ****************
```

# B

| | TaOR2 | AaOR2 | AaOR9 | TaOR10 |
|---|---|---|---|---|
| **AaOR2** | 82.67 | | | |
| **AaOR9** | 50.14 | 51.49 | | |
| **TaOR10** | 53.51 | 53.78 | 69.79 | |
| **AaOR10** | 52.85 | 52.85 | 69.79 | 78.34 |
